# Deep-learning on-chip DSLM enabling video-rate volumetric imaging of neural activities in moving biological specimens

**DOI:** 10.1101/2021.05.31.446320

**Authors:** Xiaopeng Chen, Junyu Ping, Yixuan Sun, Chengqiang Yi, Sijian Liu, Zhefeng Gong, Peng Fei

## Abstract

Volumetric imaging of dynamic signals in a large, moving, and light-scattering specimen is extremely challenging, owing to the requirement on high spatiotemporal resolution and difficulty in obtaining high-contrast signals. Here we report that through combing a microfluidic chip-enabled digital scanning light-sheet illumination strategy with deep-learning based image restoration, we can realize isotropic 3D imaging of crawling whole *Drosophila* larva on an ordinary inverted microscope at single-cell resolution and high volumetric imaging rate up to 20 Hz. Enabled with high performances even unmet by current standard light-sheet fluorescence microscopes, we *intoto* record the neural activities during the forward and backward crawling of 1^st^ instar larva, and successfully correlate the calcium spiking of motor neurons with the locomotion patterns.

## 1. Introduction

Light-sheet Fluorescence Microscopy (LSFM) allows three-dimensional imaging of live samples with high speed and low photo-bleaching.^1–7^ To push the limit of spatiotemporal performance, a variety of LSFM implementations have evolved from the classic selective plane illumination microscopy (SPIM) or digital scanning light-sheet microscopy (DSLM), to provide superior-quality imaging of samples from single cells to entire organism.^8–17^ However, unlike the light-field recording based 3D imaging,^18–22^ LSFM requires recording a sequence of 2D plane images to reconstruct a 3D volume at sufficiently-high spatial resolution, the temporal resolution is thus still fundamentally compromised by the extended acquisition time of the camera, making the clear observation of millisecond-time-scale biological dynamics, e.g., neural activities-related locomotion, across a large 3D tissues, e.g. whole larva, yet extremely difficult. Some ultrafast LSFM modality can achieve video-rate 3D imaging of dynamic samples through using high-speed opto-mechanics as well as opto-electronics to observe a confined field of view (FOV).^23–25^ Since even high-end LSFM is largely incapable of 3D dynamic imaging, the relatively simple formats of LSFM^26–29^ or cost-effective LFSM based on widely-used epifluorescence microscopes^30–32^ are even farther from this desired goal. This limit fundamentally comes from the relatively small space bandwidth product (SBP) of microscopes, and thus causes an inevitable compromise between the spatial and temporal resolution (speed). Several multi-frame-based computational approaches have been combined with LSFM to increase the spatial resolution but with compromised speed.^9,33,34^ The recurrent concern on how to improve the speed without sacrificing the accuracy is previously addressed by single-image super-resolution (SISR) approaches, such as dictionary search and compressed sensing, but only limited to certain types of spare signals.^35,36^ The recent advent of deep learning-enabled image restoration has brought big impact to the microscopy field,^22,37–41^ with the trained neural network capable of directly deducing a higher-quality image based on a single low-quality measurement. Deep learning-based restoration has been also applied to the denoising of microscopy images, in which the acquisition of qualitied microscopy data for network training is relatively difficult.^39,42^ With gaining the capability of recovering unsatisfactory signals acquired under coarse sampling and short exposure conditions, SISR-enabled LSFM could reach an improved speed without sacrificing the spatial resolution.^37–39^

Based on these state-of-the-art developments, we herein report a deep-learning on-chip (DO)-DSLM approach that combines a scanned light-sheet illumination add-on setup with efficient deep neural network (DNN)^37^, and allows a 2D inverted microscope to implement video-rate 3D imaging of dynamic processes. A specially arranged DSLM add-on in conjunction with a customized microfluidic device enables a wide-field microscope with 3D optical sectioning capability of dynamic samples. The integrated DNN-based restoration procedure further computationally improves the ∼15-µm (FWHM) ambiguous cellular resolution to ∼8-µm (FWHM) isotropic single-cell resolution, and increases the image signal-to-noise ratio (SNR) from ∼4 to ∼11. We demonstrate this DO-DSLM approach through the real-time imaging of the motor neuron activities in crawling *Drosophila* larva. As compared to very poor performance by the original epi-illumination of inverted microscope, or still fuzzy and noisy 3D reconstruction caused by the video-rate z-scan with coarse step size (∼7.5 µm), DO-DSLM hybrid strategy can readily yield 3D microscopic video of dynamic biological processes, with over 2-fold-enhanced axial resolution and notably higher SNR, allowing otherwise indistinguishable neuron calcium signals from the moving-and-scattering tissues to be super-resolved with accurate intensity also recovered. As a result of significantly improved spatiotemporal performance by this DO-DSLM approach, we further demonstrate the quantitative analysis of the neural activities and their correlations with the forward and backward locomotion of the behaving worm.

## 2. Experimental setup of DO-DSLM

We first constructed a DSLM setup on a conventional inverted microscope, to enable 3D optical sectioning of tissues. This readily-assembled setup majorly consisted of three parts (Fig. 1a, S1†a, S2† and S3†) as: (i) an existing inverted microscope (IX73, Olympus) for fluorescence detection, (ii) a self-built plane illumination module for generating a horizontal scanning laser sheet with ∼7-μm (FWHM) thickness, and (iii) a piezo-actuated mechanical module for axial scanning of the sample. A three-layer microfluidics chip (Fig. 1b and S1†b) containing control valves was designed to host live transgenic *Drosophila* larva (first instar, ventral neural cord (VNC) labelled with GFP) inside a micro chamber (Fig. 1b, 3000 × 500 μm^2^), thus allowing it freely move within the FOV of a 4X detection objective, for sustained high-speed imaging. The *Drosophila* larva is ∼200 μm in width and 700 μm in length, for which our imaging setup is sufficient for optically resolving signals of single neurons with a large FOV. It should be noted that the side wall of the microfluidic chip was optically flattened to allow the high-quality light-sheet illumination in the chip (Fig. S5†). To provide fast and low-phototoxicity 3D acquisition, a high-frequency coarse axial scanning of the crawling worm was implemented by a piezo stage at 10 Hz, correspondingly yielding 3D image stacks with ∼7.5-μm step size and at a rate of 20 volumes per second (Fig. 1c). As compared to previous obliquely-arranged light-sheet microscopes that are compatible with on-chip imaging,^25,43^ our horizontal plane illumination mode from the flatten side facet allowed the 3D image with the same volume size to be obtained at a higher speed, owing to the notable reduction of scanning range.

**Fig. 1.**
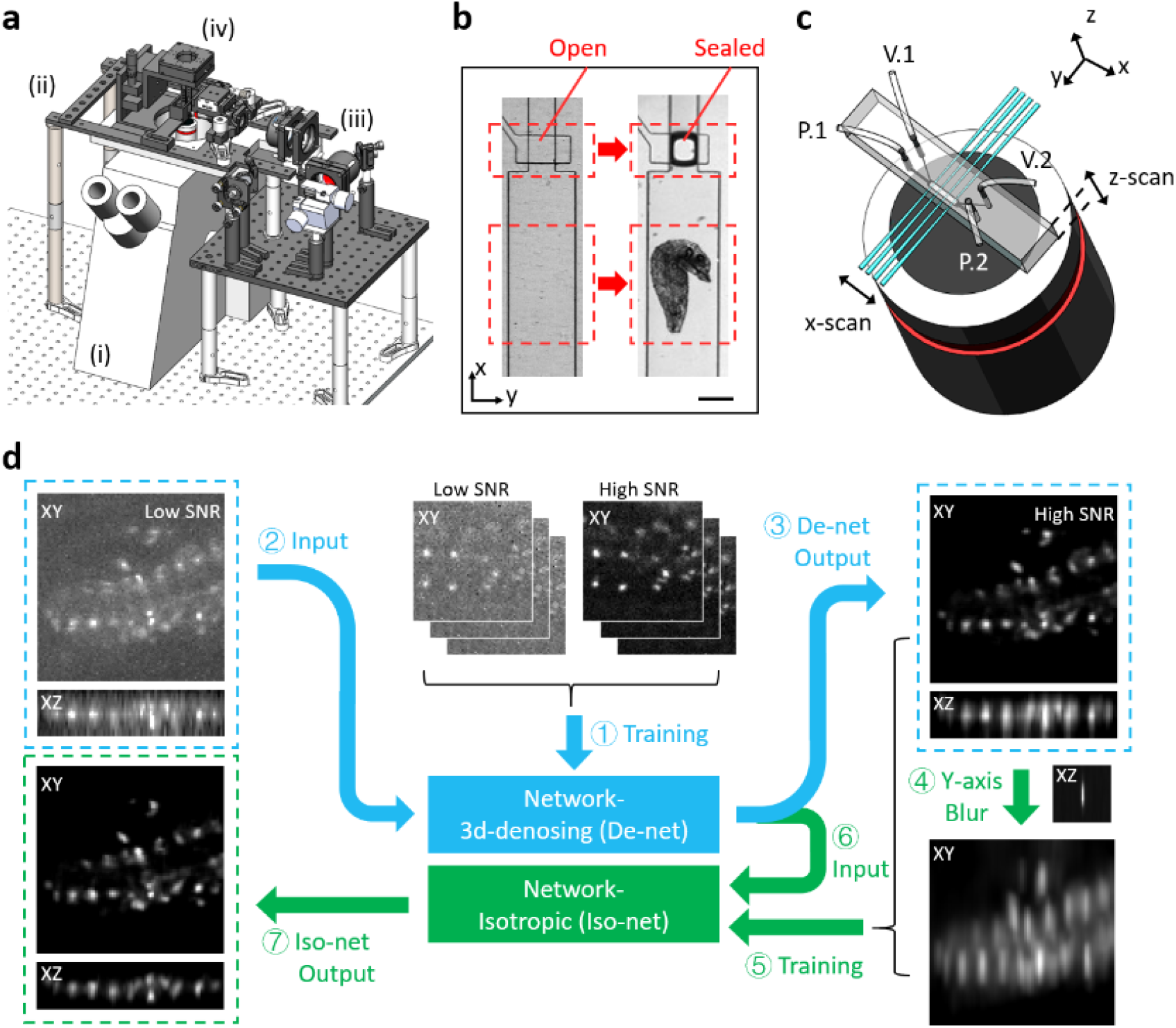
DO-DSLM procedure. (a) Design of the on-chip DSLM system with four major components indicated by (i)-(iv). (i) an inverted microscope as host (IX73, Olympus), (ii) a customized bracket above the microscope, (iii) a self-built plane illumination module, and (iv) a piezo-actuated mechanical module. (b) *Drosophila* under control in a three-layer microfluidics chip. The valve keeps open before a larva being loaded into the chamber (left). Then it keeps sealed once the larva appears in the FOV (right). Scale bar. 250 μm. (c) Geometry of the on-chip DSLM at working status. The scanned light-sheet optically sections the chamber through PDMS from the optically-flattened facet. The light sheet keeps stationary at the vicinity of the focal plane of the detection objective. The microfluidics chamber is moved across the light-sheet for a distance of ∼60 μm along z direction at 10 Hz, realizing 20 volumetric scanning per second. (d) Deep-learning image restoration procedure. (1) De-net training. (2) Low-SNR raw images input. (3) High-SNR images output (De-net Output). (4) Lateral blurring using system PSF. (5) Iso-net training. (6) Low axial resolution images input. (7) High axial resolution images output (Iso-net Output).

The raw on-chip DSLM images with compromised axial resolution and SNR induced by the coarse scanning, short exposure and tissue scattering were further restored by deep-learning-based denoising and isotropic resolution enhancement (Fig. 1d and S6†). Fig. 1d shows the schematic of the denoised- and-isotropic network (De-iso-Net). We acquired 3D stacks of static *Drosophila* larvae with different exposure time (10 ms and, 100 ms) for denoising sub-network (De-net) training (step 1). Then the input raw images were restored by the trained De-net (step 2) to generate high-SNR but anisotropic outputs (step 3), which were further degraded to generate synthetic low-resolution axial slices (step 4) for isotropic sub-network (Iso-net) training (step 5). Finally, the denoised output of De-net was resliced into a stack of axial slices (step 6) and restored by trained Iso-net to generate final denoised and isotropic results (step 7).

## 3. Results and discussion

### 3.1. 3D imaging and restoration of fluorescent bead and static *Drosophila* on a chip

With optically-flatten side wall that led to smooth air-PDMS interface, the horizontal laser sheet covered the full width of chamber to illuminate the entire sample at any time point, Also, unlike 45-degree light-sheet microscopes that need to scan the full width of the chamber (500 μm in our experiment), DO-DSLM only scans the height of the chamber (up to 100 μm in our experiment), notably improving the acquisition speed through the reduction of scanning range. Therefore, the system could exhibit its full spatiotemporal performance for imaging dynamic samples. We first measured the point-spread-function (PSF) of DO-DSLM by imaging fluorescent beads (1-μm diameter, Lumisphere, BaseLine Chromtech) packed inside the chip (5-μm step size). As shown in Fig. 2a, both De-Net and De-iso-net notably improve the SNR of beads obtained under a short exposure time of 5 ms. Meanwhile, they both keep a high structural similarity (SSIM) with the high-SNR ground truth acquired under 100-ms exposure time. In addition, the De-iso-net further provides isotropic resolution which could better resolve the single neurons in three dimensions. As shown in Fig. 2b and c, the axial PSF extent was improved from ununiform 5-12 μm (FWHM value) in the raw and De-net results to uniform 5 μm in De-iso-net results.

**Fig. 2.**
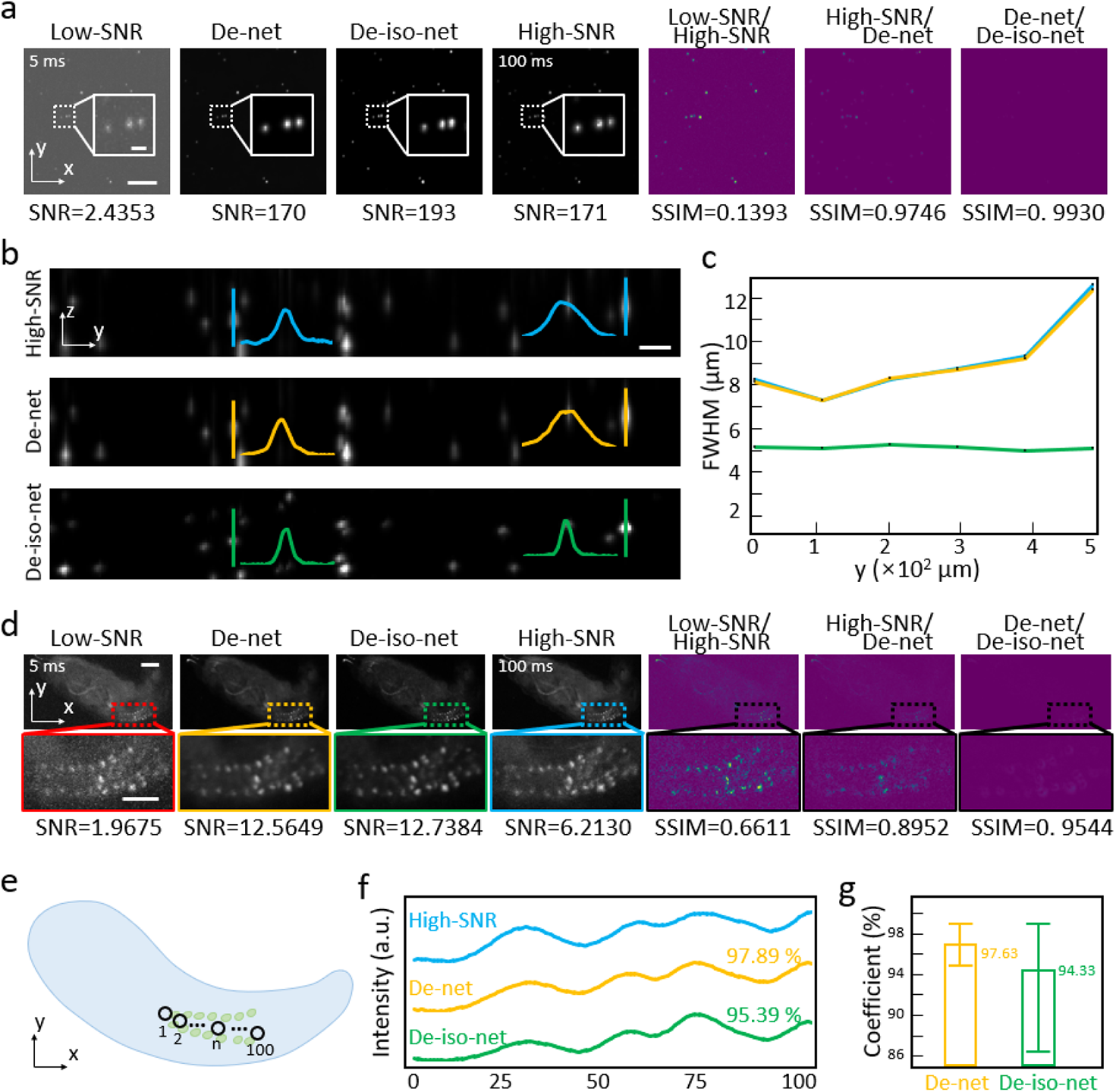
Performance of DO-DSLM. (a) Maximum intensity projections (MIPs) of x-y planes of fluorescent beads (500 μm × 500 μm). From left to right are, low signal-to-noise (SNR) image captured with an exposure time of 5 ms (Low-SNR), restoration of Low-SNR by De-net, restoration of De-net data by Iso-net, high SNR image captured with an exposure time of 100 ms in situ (High SNR), error map of low-SNR and high-SNR, error map of high-SNR and De-net, error map of De-net and De-iso-net, respectively. Scale bar: 100 μm. Inset scale bar: 10 μm. (b) MIPs of y-z planes of fluorescent beads (500 μm × 80 μm). From top to bottom are, high-SNR results, De-net results and De-iso-net results, respectively. Scale bar: 20 μm. (c) The variation of lateral full width half maximums (FWHMs) of resolved beads along the 500-μm FOV, comparing the axial resolution of LR, De-net and De-iso-net results. (d) MIPs of x-y planes of a static *Drosophila* larva. From left to right are, low signal-to-noise (SNR) image captured with an exposure time of 5 ms (Low-SNR), restoration of Low-SNR by De-net, restoration of De-net data by Iso-net, high SNR image captured with an exposure time of 100 ms in situ (High-SNR), error map of low-SNR and high-SNR, error map of high-SNR and De-net, error map of De-net and De-iso-net, respectively. Scale bar: 100 μm. Inset scale bar: 50 μm. (e) Schematic of a *Drosophila* larva with 100 small round regions indicated. Spot size: 4 pixels in diameter. (f) The normalized intensity along 100 samples selected in (e), comparing the correlation of high-SNR, De-net and De-iso-net. (g) Statistics of the correlation coefficient of De-net and De-iso-net.

We further validated the DO-DSLM imaging performance on a 1st instar Drosophila larva (5-μm step size). Both De-Net and De-iso-net can improve the SNR of neuron signals ∼6 folds from ∼2 (under 5-ms exposure) to ∼12, a value even higher than the signal SNR of ∼6 obtained under 100 ms exposure (Fig. 2d). We further randomly sampled 100 small regions (size 4 pixels) from De-net and De-iso-net results (Fig. 2e), and compared their average intensity profiles with those from high-SNR ground-truth data (Fig. 2f). The averaged coefficients are 97% for De-net and 94% for De-iso-net (Fig. 2g), verifying the high intensity accuracy in reconstructed signals which is sufficient for analysing the neural activities.

### 3.2. Demonstration of the necessity of high volumetric rate and networks

The crawling speed of a freely-moving *Drosophila* larva can be over 100 μm per second. Thus, it is essential to impletement 3D imaging at high volume rate to capture the calcium signals of single neuron moving and blinking in three dimensions. We showed that the moving larva captured at 4 volumes per second (vps, 20 ms exposure, ∼7.5-μm step size, 8 frames per volume) has suffered from severe motion blurring in the maximum-intensity-projections (MIPs) of x-y plane, causing most of neurons being undistinguishable. Meanwhile, in the reconstructed x-z plane, the signal distoration also made the follow-up signal analysis extremely difficult (Fig. 3a, left). On the contrary, when imaging at a volumetric rate of 20 Hz by a short exposure time and large z step, though the motion blurring and distoration almost vanished, the results showed the issue of low SNR instead (Fig. 3a, middle). Combined with the ambiguous axial resolution, it also caused big difficulty to the signal analysis. Then, the De-iso-net was applied to recover the image SNR and insufficient axial resolution for the 20-vps data, sucessfully achieving 3D signal reconstruction at satisifying single-neuron resolution and 20 Hz high speed. While both the 4-vps and raw 20-vps DSLM data are inadequte to completely reveal the nerual activities in a live moving larva, De-iso-net-enhanced DLSM imaging substantially addresses this challenge. Meanwhile, since the signal intensities before and after the network reconstruction are highly correlated, DO-DSLM has been proved to be suited for high-speed 3D quantitative imaging of biological dynamics. As we superposed the MIPs of ten consecutive DO-DSLM volumes with each volume represented by one color (Fig. 3b-c), we clearly visualized the tracks of 2 adjacent neurons (N.1 and N.2) with also quantifying their calcium intensity variations during the motion.

**Fig. 3.**
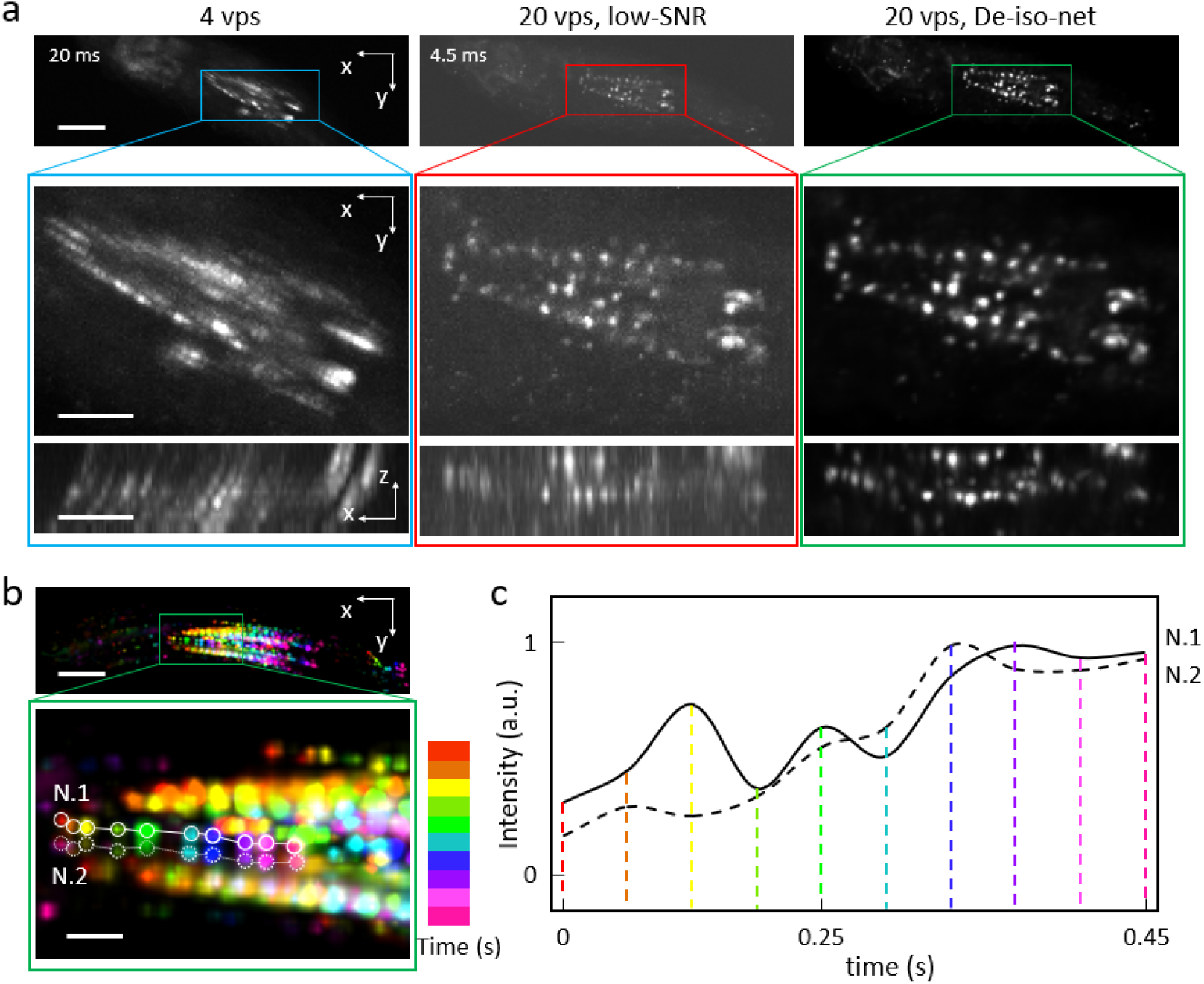
DO-DSLM imaging of freely-moving *Drosophila* larva with different volumetric imaging rates. (a) MIPs of x-y planes of freely-moving *Drosophila* larvae. From left to right are, larva captured at 4 vps, 20 vps and the corresponding De-iso-net results of 20 vps. Insets show the magnification of neuron regions at x-y planes and x-z planes. Scale bar: 100 μm. Inset scale bar: 50 μm. (b) Superposition of MIPs of x-y planes of ten consecutive volumes (De-iso-net results, 0.5 s in total). Color represents the different volumes. Two neurons are traced (N.1 & N.2) over a 0.5 s period, showing the significant signal drifts even at sub-second timescale. Scale bar:100 μm. Inset scale bar: 20 μm. (c) Normalized intensity variation of N.1 and N.2. during 0.5 s.

### 3.3. Neural activities of freely moving *Drosophila* larva

We further validated the performance of DO-DSLM through restoring a time-lapse sequence of 100 3D image stacks acquired in a 5-second live imaging of free-moving *Drosophila* larva. As the schematic shown (Fig. 4a), the raw data is noisy (first column). After the processing of De-net, it looks clearer (second column). Finally, after the processing of Iso-net, we achieve isotropic resolution in three dimensions (third column). We show the details of three volume (t=0, t=2, and t=4 s) as examples for proving the effective and significance of the networks in dealing with raw data of a freely moving *Drosophila* larva (Fig. 4b). While the average SNR has been improved from ∼4 to ∼11 by De-net, the axial elongation of neurons still made the neuron identification (dots in Imaris) difficult. After the further processing by De-Iso-net, the neurons could be accurately quantified by a spot in Imaris and their SNR remained unchanged.

**Fig. 4.**
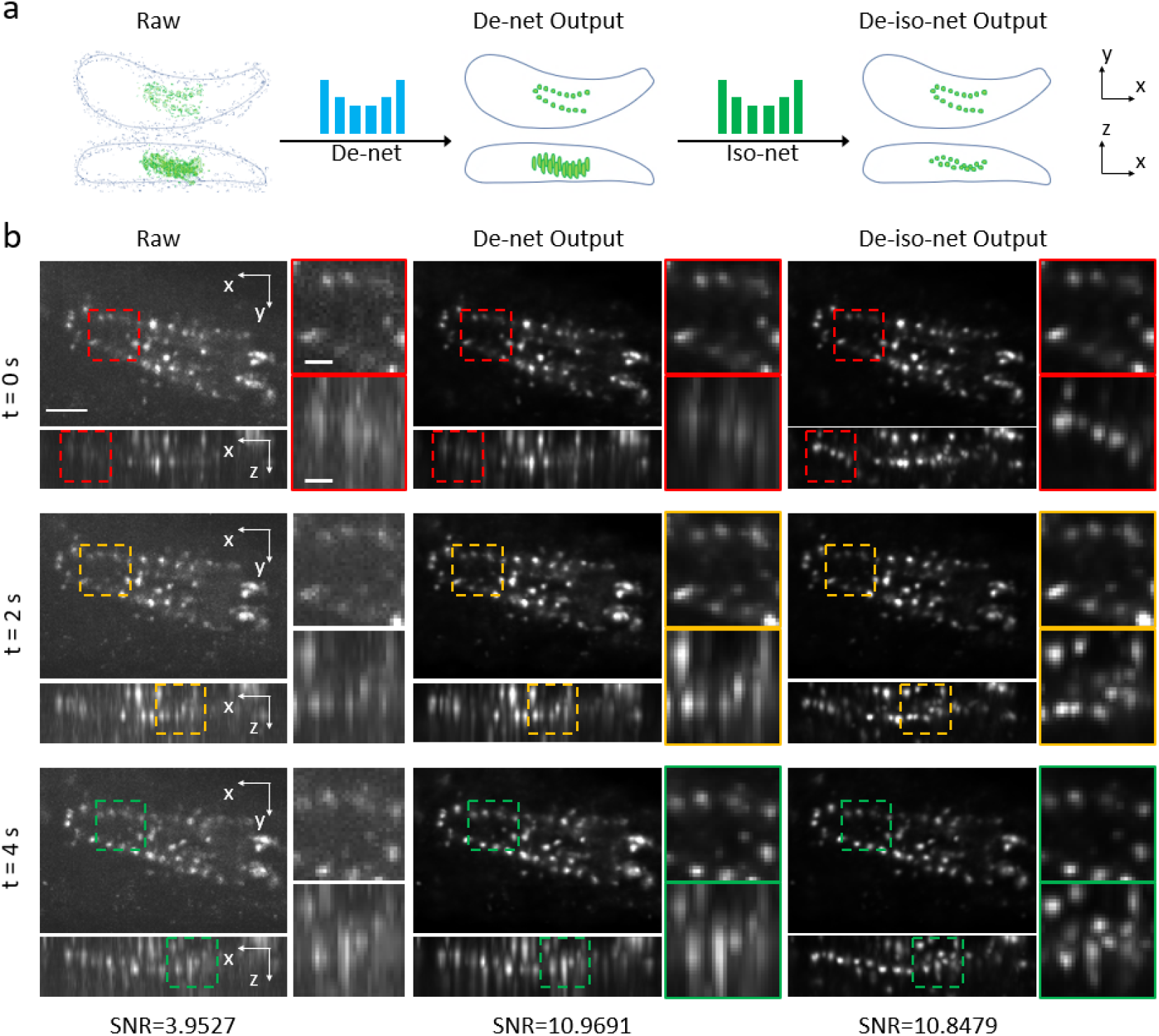
De-iso-net restoration of a freely-moving *Drosophila* larva at 20-Hz volumetric imaging rate. (a) Schematic of *Drosophila* signal restoration which contains progressive denoising and resolution enhancement. (b) Restoration of three consecutive volumes (at t=0 s, t=2 s and t=4 s) of *Drosophila* larva with VNCs labelled by GFP. The first column shows the noisy neurons in x-y and x-z plane. The second column shows the restoration output of De-net with respect to the first column. The third column shows the final restoration output of De-iso-net correspondingly. Scale bar: 50 μm. Inset scale bar: 10 μm.

### 3.4. Locomotion and neural activities of freely-moving *Drosophila* larvae

As compared to SCAPE microscopy, which previously achieved a volumetric imaging rate of 11 vps for *Drosophila* larva, our simple inverted microscope-based DO-DSLM is now able to implement isotropic 3D imaging of VNC motor neurons at deeper tissues, with wider FOV and at higher volume rate (20 Hz or even higher). We analysed the high-SNR and isotropic data reconstructed by De-iso-net, revealing the correlation between the locomotion and neural activities during larvae movement (Fig. 5, Video S1). Due to the intrinsic symmetrical structure of the motor neurons, we only need to trace one side of the VNCs. Trough analysing the sequential DO-DSLM image stacks, we visualized the traces of 8 motor neurons (neurons are color-coded, N.1 to N.8 is in order of the larval segment) during the forward and backward crawling of the worms, as shown in Fig. 5a and d, respectively. The magnified views in Fig. 5b and 5e further detailed the neuron traces, for example, the neuron N.1 (red) travelling through the red track from the right to the left during a 4-second period (Fig. 5b), which corresponded to a forward moving larva whose anterior part is towards the left. Furthermore, the motor neural activities and their association with the larval locomotion patterns could be also quantitatively revealed, as shown in Fig. 5c, f. A cycle of crawling wave was usually accompanied with large displacement of larva body. Therefore, it is reasonable to analyse the larvae’s locomotion through tracking the speeds of neurons. As indicated by the black arrows in Fig. 5c and f, we identified two and three velocity peaks, which corresponded to two and three cycles of crawling waves during the observation periods. These identified crawling waves also shows obvious correlations with the sequential calcium signalling of VNC neurons, as shown by the dash lines, indicating that the rises and falls (from N.8 to N.1, tail to head) of fluorescence-indicated neural activities in VNC neurons are consistent with the larval forward/ backward crawling.

**Fig. 5.**
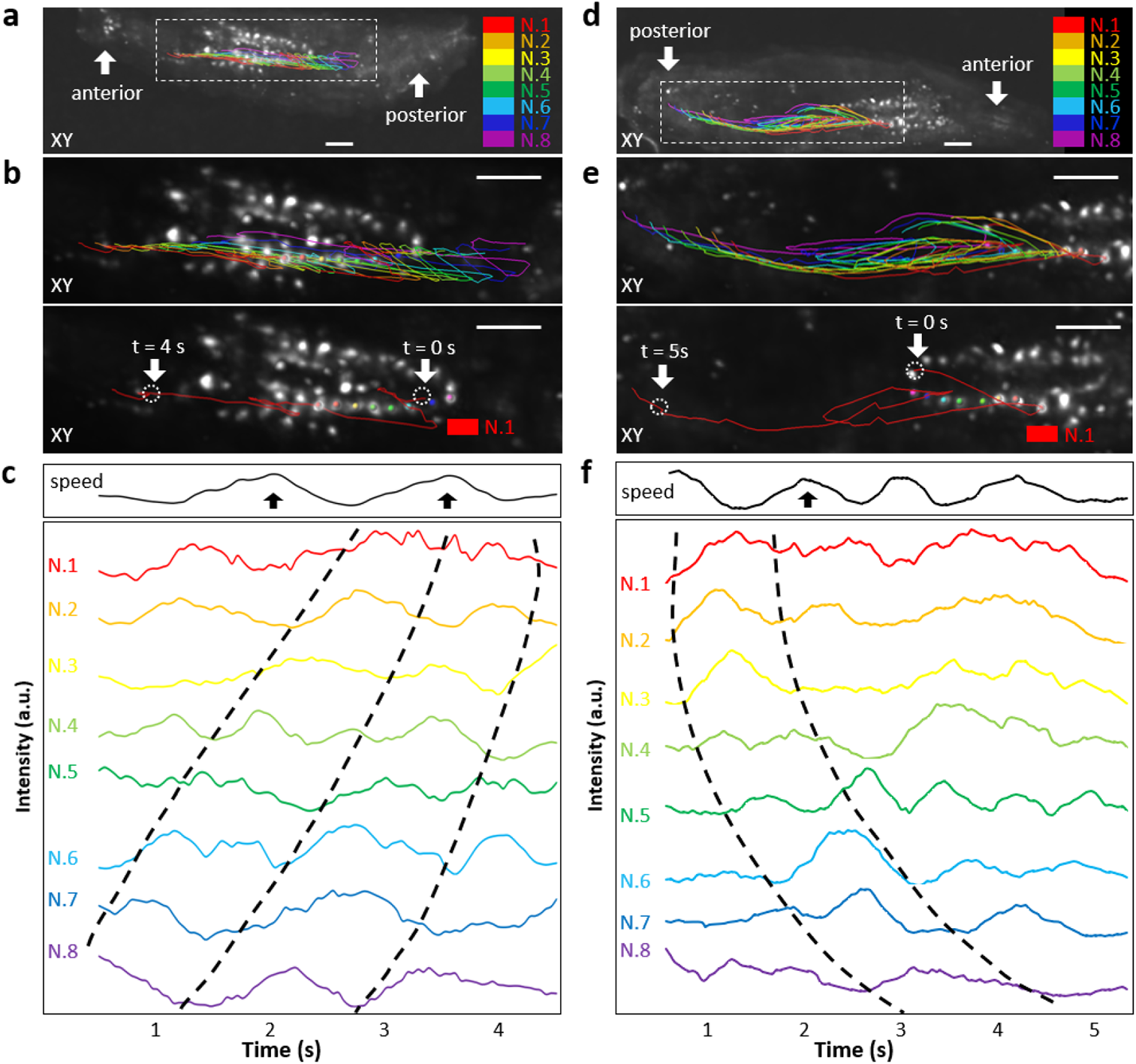
Motion analysis and neurons calcium measurement for forward and backward crawling larvae. (a) MIP of x-y plane of a forward crawling larva. The anterior is to the left and the posterior is to the right. 8 neurons and their tracks (N.1∼N.8) are labelled with different colors. Scale bar: 50 μm. (b) Magnification of the neuron regions selected in (a) by the dotted rectangular box (top). A separate neuron (N.1) and its track (red line) are shown (bottom). Track duration: 4.5 s. Scale bar: 50 μm. (c) Neuron speed and corresponding calcium intensity variation during 4.5 s. Two black arrows indicate the large displacement of the larva (top, speed variation curve). Three dotted lines indicates the calcium signal transmission from N,8 to N,.1 in proper order (bottom, calcium intensity curves). The (d), (e), (f) show similar results for a backward crawling larva.

## 4. Conclusions

Through the win-win combination of DSLM illumination that enables conventional inverted microscope with strong optical sectioning capability, microfluidic chip that is compatible with light-sheet illumination for efficient sample control / imaging, and deep-learning-based restoration that notably improves the image quality, our DO-DSLM strategy achieved 20-Hz video-rate 3D functional imaging of behaving *Drosophila* larva at single-cell resolution. The up-limit of the volumetric imaging rate of DO-DSLM can reach 50-Hz with using faster sCMOS camera and *Drosophila* sample containing stronger labelling. As a result of the superior imaging, we quantitatively analysed the neural activities in acting *Drosophila* larvae and revealed their correlations with forward and backward crawling locomotion behaviours. DO-DSLM shows how a high-quality 3D dynamic imaging can be realized efficiently and cost-effectively in a laboratory which already has a common fluorescence microscope. Thus, we expect this approach can be widely applied to different biomedical applications under various resource-limited environments.

## Author Contributions

P. F. conceived the idea. P. F. and G. Z. oversaw the project. X. C., J. P., Y.S., S. L. designed and implemented the experiments:, Y. S. and Z. G. provided resources and advice on biology. X. C., C. Y., J. P. conducted the image reconstruction and analysis. P. F., X. C., J. P., Z. G. wrote the manuscript.

## Acknowledgements

This research has received the funding support from the National Natural Science Foundation of China (21874052, 21927802), the ZHEJIANG LAB (111002-AC002). The authors would like to thank Zhaofei Wang, Tingting Zhu, Jinrun Zhou for their helpful comments and discussion.

## References

1 J. Huisken, J. Swoger, F. Del Bene, J. Wittbrodt and E. H. K. Stelzer, Science, 2004, 305, 1007–1009.

2 J. Huisken and D. Y. R. Stainier, Opt. Lett., 2007, 32, 2608–2610

3 Y. Wu, A. Ghitani, R. Christensen, A. Santella, Z. Du, G. Rondeau, Z. Bao, D. Colón-Ramos and H. Shroff, Proc. Natl. Acad. Sci. U. S. A., 2011, 108, 17708–17713.

4 Y. Wu, P. Wawrzusin, J. Senseney, R. S. Fischer, R. Christensen, A. Santella, A. G. York, P. W. Winter, C. M. Waterman, Z. Bao, D. A. Colón-Ramos, M. McAuliffe and H. Shroff, Nat. Biotechnol., 2013, 31, 1032–1038.

5 H. U. Dodt, U. Leischner, A. Schierloh, N. Jährling, C. P. Mauch, K. Deininger, J. M. Deussing, M. Eder, W. Zieglgänsberger and K. Becker, Nat. Methods, 2007, 4, 331–336.

6 M. Mickoleit, B. Schmid, M. Weber, F. O. Fahrbach, S. Hombach, S. Reischauer and J. Huisken, Nat. Methods, 2014, 11, 919–922.

7 P. J. Keller, A. D. Schmidt, J. Wittbrodt and E. H. K. Stelzer, Science, 2008, 322, 1065–1069.

8 T. A. Planchon, L. Gao, D. E. Milkie, M. W. Davidson, J. A. Galbraith, C. G. Galbraith and E. Betzig, Nat. Methods, 2011, 8, 417–423.

9 P. J. Keller, A. D. Schmidt, A. Santella, K. Khairy, Z. Bao, J. Wittbrodt and E. H. K. Stelzer, Nat. Methods, 2010, 7, 637–642.

10 K. M. Dean, P. Roudot, E. S. Welf, G. Danuser and R. Fiolka, Biophys. J., 2015, 108, 2807–2815.

11 R. McGorty, H. Liu, D. Kamiyama, Z. Dong, S. Guo and B. Huang, Opt. Express, 2015, 23, 16142–16153.

12 L. Gao, L. Shao, C. D. Higgins, J. S. Poulton, M. Peifer, M. W. Davidson, X. Wu, B. Goldstein and E. Betzig, Cell, 2012, 151, 1370–1385.

13 F. Cella Zanacchi, Z. Lavagnino, M. Perrone Donnorso, A. Del Bue, L. Furia, M. Faretta and A. Diaspro, Nat. Methods, 2011, 8, 1047– 1050.

14 M. Hagiwara, F. Peng and C. M. Ho, Sci. Rep., 2015, 5, 1–7.

15 P. Theer, D. Dragneva and M. Knop, Sci. Rep., 2016, 6, 1–9.

16 T. Vettenburg, H. I. C. Dalgarno, J. Nylk, C. Coll-Lladó, D. E. K. Ferrier, T. Čižmár, F.J. Gunn-Moore and K. Dholakia, Nat. Methods, 2014, 11, 541–544.

17 J. C. M. Gebhardt, D. M. Suter, R. Roy, Z. W. Zhao, A. R. Chapman, S. Basu, T. Maniatis and X. S. Xie, Nat. Methods, 2013, 10, 421– 426.

18 R. Prevedel, Y. G. Yoon, M. Hoffmann, N. Pak, G. Wetzstein, S. Kato, T. Schrödel, R. Raskar, M. Zimmer, E. S. Boyden and A. Vaziri, Nat. Methods, 2014, 11, 727–730.

19 N. C. Pégard, H.-Y. Liu, N. Antipa, M. Gerlock, H. Adesnik and L. Waller, Optica, 2016, 3, 517–524.

20 L. Cong, Z. Wang, Y. Chai, W. Hang, C. Shang, W. Yang, L. Bai, J. Du, K. Wang and Q. Wen, Elife, 2017, 6, 1–20.

21 N. Wagner, N. Norlin, J. Gierten, G. de Medeiros, B. Balázs, J. Wittbrodt, L. Hufnagel and R. Prevedel, Nat. Methods, 2019, 16, 497–500.

22 Z. Wang, L. Zhu, H. Zhang, G. Li, C. Yi, Y. Li, Y. Yang, Y. Ding, M. Zhen, S. Gao, T. K. Hsiai and P. Fei, Nat. Methods, 2021, 18, 551– 556.

23 F. O. Fahrbach, F. F. Voigt, B. Schmid, F. Helmchen and J. Huisken, Opt. Express, 2013, 21, 21010–21026.

24 M. B. Bouchard, V. Voleti, C. S. Mendes, C. Lacefield, W. B. Grueber, R. S. Mann, R. M. Bruno and E. M. C. Hillman, Nat. Photonics, 2015, 9, 113–119.

25 R. D. Vaadia, W. Li, V. Voleti, A. Singhania, E. M. C. Hillman and W. B. Grueber, Curr. Biol., 2019, 29, 935-944.e4.

26 M. Stefaniuk, E. J. Gualda, M. Pawlowska, D. Legutko, P. Matryba, P. Koza, W. Konopka, D. Owczarek, M. Wawrzyniak, P. Loza-Alvarez and L. Kaczmarek, Sci. Rep., 2016, 6, 1–9.

27 Z. Yang, P. Haslehurst, S. Scott, N. Emptage and K. Dholakia, Sci. Rep., 2016, 6, 1–7.

28 P. Liao, M. Jiang, Z. Chen, F. Zhang, Y. Sun, J. Nie, M. Du, J. Wang, P. Fei and Y. Huang, Proc. Natl. Acad. Sci. U. S. A., 2020, 117, 25628–25633.

29 P. G. Pitrone, J. Schindelin, L. Stuyvenberg, S. Preibisch, M. Weber, K. W. Eliceiri, J. Huisken and P. Tomancak, Nat. Methods, 2013, 10, 598–599.

30 Z. Guan, J. Lee, H. Jiang, S. Dong, N. Jen, T. Hsiai, C.-M. Ho and P. Fei, Biomed. Opt. Express, 2016, 7, 194–208.

31 F. Zhao, L. Zhu, C. Fang, T. Yu, D. Zhu and P. Fei, Biomed. Opt. Express, 2020, 11, 7273–7285.

32 F. Zhao, Y. Yang, Y. Li, H. Jiang and X. Xie, J. Biophotonics, 2020, 13, 1–15.

33 J. Nie, S. Liu, T. Yu, Y. Li, J. Ping, P. Wan, F. Zhao, Y. Huang, W. Mei, S. Zeng, D. Zhu and P. Fei, Adv. Sci., 2019, 7, 1-11.

34 B. C. Chen, W. R. Legant, K. Wang, L. Shao, D. E. Milkie, M. W. Davidson, C. Janetopoulos, X. S. Wu, J. A. Hammer, Z. Liu, B. P. English, Y. Mimori-Kiyosue, D. P. Romero, A. T. Ritter, J. Lippincott-Schwartz, L. Fritz-Laylin, R. D. Mullins, D. M. Mitchell, J. N. Bembenek, A. C. Reymann, R. Böhme, S. W. Grill, J. T. Wang, G. Seydoux, U. S. Tulu, D. P. Kiehart and E. Betzig, Science, 2014, 346, 1-12.

35 C. Fang, T. Yu, T. Chu, W. Feng, F. Zhao, X. Wang, Y. Huang, Y. Li, P. Wan, W. Mei, D. Zhu and P. Fei, Nat. Commun., 2021, 12, 1– 13.

36 L. Zhu, W. Zhang, D. Elnatan and B. Huang, Nat. Methods, 2012, 9, 721–723.

37 M. Weigert, U. Schmidt, T. Boothe, A. Müller, A. Dibrov, A. Jain, B. Wilhelm, D. Schmidt, C. Broaddus, S. Culley, M. Rocha-Martins, F. Segovia-Miranda, C. Norden, R. Henriques, M. Zerial, M. Solimena, J. Rink, P. Tomancak, L. Royer, F. Jug and E. W. Myers, Nat. Methods, 2018, 15, 1090–1097.

38 Z. Hao, C. H. F. Ang, X. I. X. Ie, Y. I. Y. Ang, W. E. I. M. Ei, D. I. J. In and P. E. N. G. F. Ei, Biomed. Opt. Express, 2019, 10, 1044–1063.

39 Z. Hao, Y. Zhao, C. Fang, G. Li, M. Zhang, Y. Zhang and P. Fei, Optica, 2020, 7, 1627–1640.

40 Y. Wu, Y. Rivenson, H. Wang, Y. Luo, E. Ben-David, L. A. Bentolila, C. Pritz and A. Ozcan, Nat. Methods, 2019, 16, 1323–1331.

41 W. Ouyang, A. Aristov, M. Lelek, X. Hao and C. Zimmer, Nat. Biotechnol., 2018, 36, 460–468.

42 X. Le, C. Fang, L. Zhu, Y. Wang, T. Yu, Y. Zhao, D. Zhu and P. Fei, Opt. Express, 2020, 28, 30234–30247.

43 X. Han, Y. Su, H. White, K. M. O’Neill, N. Y. Morgan, R. Christensen, D. Potarazu, H. D. Vishwasrao, S. Xu, Y. Sun, S. Huang, M. W. Moyle, Q. Dai, Y. Pommier, E. Giniger, D. R. Albrecht, R. Probst and H. Shroff, Lab Chip, 2021, 21, 1549–1562.

