## Supplementary Material for "Deep-learning on-chip DSLM enabling video-rate volumetric imaging of neural activities in moving biological specimens"

### Supplementary Information

#### 1. Methods

##### 1.1 Strains

*Drosophila melanogaster* strains were maintained on standard culture medium at 25°C, 60% humidity. The larva carried a R36G02-GAL4 transgene (Pfeiffer, B.D., Jenett, A., Hammonds, A.S., Ngo, T.T., Misra, S., Murphy, C., Scully, A., Carlson, J.W., Wan, K.H., Laverty, T.R., Mungall, C., Svirskas, R., Kadonaga, J.T., Doe, C.Q., Eisen, M.B., Celniker, S.E., Rubin, G.M. (2011.2.9). GAL4 Driver Collection of Rubin Laboratory at Janelia Farm. https://flybase.org/reports/FBrf0213040) and a UAS-GCaMP6m transgene to express GCaMP6m, a calcium concentration indicator that marks cell activity, in neurons in ventral nerve cord. Calcium imaging experiments were performed with first-instar larvae to minimize scattering.

##### 1.2 DSLM illumination setup

A focused Gaussian laser beam (488 nm, CNI laser) was scanned into a thin sheet by a dual-axis galvo scanner and thereby provided sharp optical sectioning of the sample (Fig. S1a). The planar scanned fields (dotted lines) generated by the scan lenses (SL) were reimaged onto the sample plane by the tube lenses (TTL) and illumination objective (IO) (UPLXAPO4X). The axial FWHM of the beam waist on sample plane was ~7 µm. The corresponding confocal range was ~650 µm, which is enough for covering a microfluidics chamber with ~500 µm width.

##### 1.3 System design by Solidwork

The calcium fluorescence signals were collected by a commercial inverted microscope (Olympus, IX73, 4X/0.28 objective) mounted with a sCMOS camera (ORCA-Flash4.0LT). A customized bracket above the 2-axes microscope stage was set up for mounting the water chamber, and the illumination objective (Fig. S2a). An illumination module with a dual-axis galvo scanner (Cambridge, MicroMax 671 series) was placed on the right of the microscope to generate and scan the light (Fig. S2b). Finally, a sample holder was mounted on the mechanical stage (Fig. S2c). This sample modulus was used to clamp the microfluidics chip and achieve rapid scan vertically using a piezo stage (COREMORROW, P12.Z200S). All the customized parts were designed using SolidWorks and then made by CNC machining.

##### 1.4 Signal control by LabVIEW

We made the control program using Labview. The Galvo scanner and piezo stage required analog signal inputs, while the camera required digital signal input (Fig. S4). We used one sequence of triangle wave (seq 1, 480 Hz) to control the galvo scanner and another sequence (seq 2, 10 Hz) to control the piezo stage. The trigger mode of the camera was set to External start trigger and received a sequence of high-frequency digital square wave (seq 3, 160 Hz). Moreover, to achieve precise high-speed three-dimensional imaging, sequence 2 and 3 were synchronized by a DAQ device (NI PCIe 6251).

#### 1.5 Microfluidics chip fabrication, control and side-facet treatment

The Microfluidics chip was fabricated using PDMS elastomer (Sylgard 184, Dow Corning). The chip contained three layers, as shown in Fig. S1b. During experiment, the Larva was injected into the fluid layer (first layer, 5:1) by water, and then locked into the chamber by the control layer (second layer, 20:1). The third layer (10:1) simply served as the substrate. We used a syringe pump (S.0) for the larva loading, and other two syringe pumps (S.1 & S.2) for valves control. Prior to the imaging, we loaded a larva into the chamber through the Port 1 (P.1), with V.1, V.2 being off. When the larva was gently loaded into the imaging chamber, V.1 and V.2 were activated by S.1 and 2 to lock the worm inside the chamber for imaging. After that, the V.1 and V.2 were released again to allow the larva being flushed away from the imaging chamber. Finally, this cycle was applied to the second worm. It should be noted that, for high-quality illumination from the side of the chip. The usually coarse side-facets of the chip were optically flattened (Fig. S5). This was accomplished by simply attaching the side-facet to the silicon wafer spin-coated with liquid PDMS. After baking, the chip was peeled off from the wafer and the side-facet became optically smooth (Fig. S5c).

#### 1.6 Image acquisition

The height of the chamber is approximately 100 µm. The z-step of DSLM imaging was 7.5 µm with each volume contains 8 slices of images. The exposure time of the camera was 4.5 ms and the frame rate was 160 (2048×256 pixels). Meanwhile, the piezo moved the sample at a rate of 10 cycles per second, thus achieving a 20-Hz volumetric imaging rate. The exposure time for the low-SNR and high-SNR data obtained for the network training was 10 ms and 100 ms, respectively.

#### 1.7 Deep learning restoration

The raw data was restored by a 3D-denoising model (De-net) combined with an isoCARE model (Iso-net). The low-SNR data and high-SNR data were paired and trained by De-net. After training, De-net predicted high-SNR images from low-SNR three-dimensional input, which was captured at high speed by piezo stage. Later on, Iso-net convolved and down-sampled (downsampling rate 4.6) the high-SNR outputs using a simulated PSF along x or y lateral dimension, to generate the semi-synthetic data which simulated the anisotropic axial stacks. The well-trained Iso-net further improved the axial resolution of De-net's outputs and finally provided high-SNR isotropic results qualified for quantitative analysis.

The patch size of training datasets for De-net was 256×128 pixels ×16 slices, with totally 7 pairs trained. The De-net model training took less than an hour (epoch = 400) using a single Nvidia 1080 Ti GPU. The De-net model inference was fast with taking ~10 minutes to reconstruct 100 high-SNR image stacks (2048×256 × 8 voxels each). The Iso-net model training took ~ one hour (epoch = 400) with using the same GPU. Its inference took ~20 minutes to reconstruct 100 image stacks (2048×256 pixels × 37 voxels each).

#### 1.8 Data analysis

VNC neurons identification and track trace were implemented by Imaris software (Bitplane, Oxford Instrument). The neurons were semi-automatically identified using spherical regions of interest (6-µm diameter) with their intensities being traced over time. Finally, we plotted the mean intensities of the neuron calcium signals versus time F(t). We applied a normalization to the signals with formula (F(t)-min)/(max-min), where the term max denoted the maximum value of F(t) and min denoted the minimum one. The locomotion of larva was characterized by the calculated speed of the VNC neurons. The SNR was calculated by formula (A-B)/C, where A denoted the mean value of the signal region, B denoted the mean value of the background region, and C denoted the standard deviation of the background region[1]. The error map was calculated by a FIJI plugin termed 'NanoJ-SQUIRREL'. The correlation coefficient between two images was calculated by MATLAB built-in function termed 'corrcoef'.

#### 2. Supplementary Figures

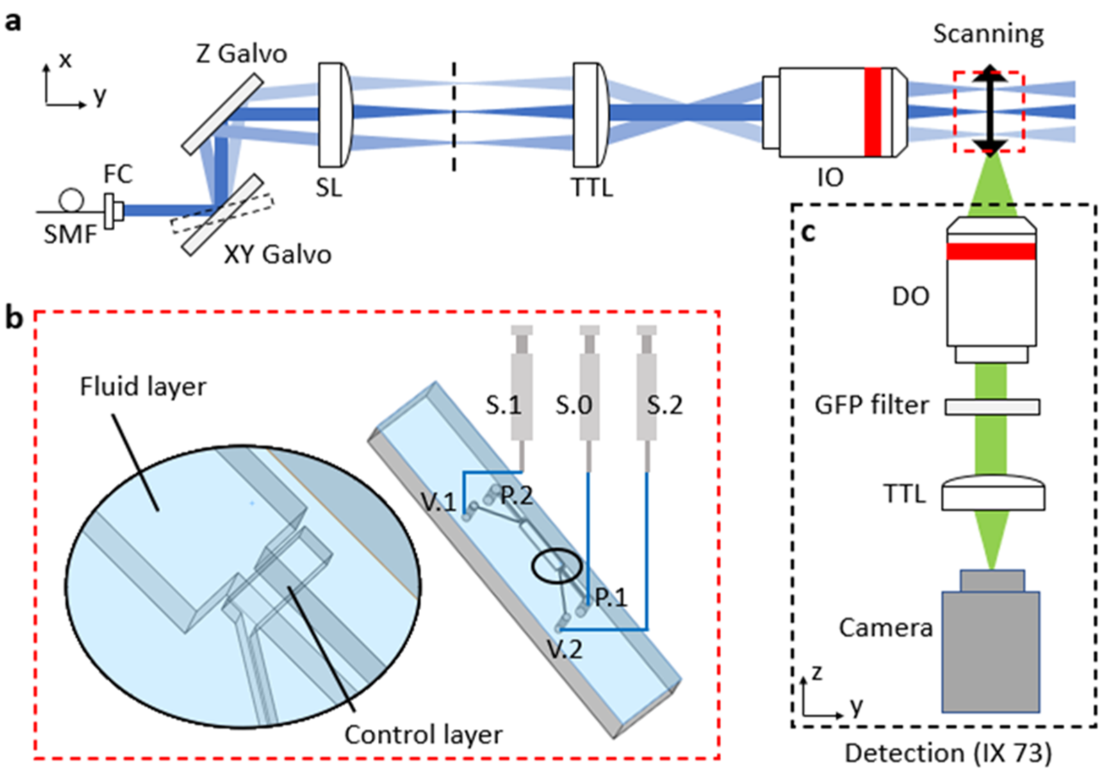


**Figure S1. optical setup and microfluidics control of DO-DSLM.** (a) Schematic of the DSLM illumination design shown in x-y view. (b) Schematic of the three-layers microfluidics chip (red dotted rectangular box in a). The Larva was loaded into the chamber through P.1 (the inlet of the microfluidics chip). Meanwhile the water flowed away from the chamber through P.2 (outlet of the microfluidics chip). S.1 and S.2 are two microfluidics valves for fluid and larva control. The inlet and outlet of the fluid layer are covered by the control layer valves, as shown in the inset. (c) 2D schematic of the detection design shown in y-z view.

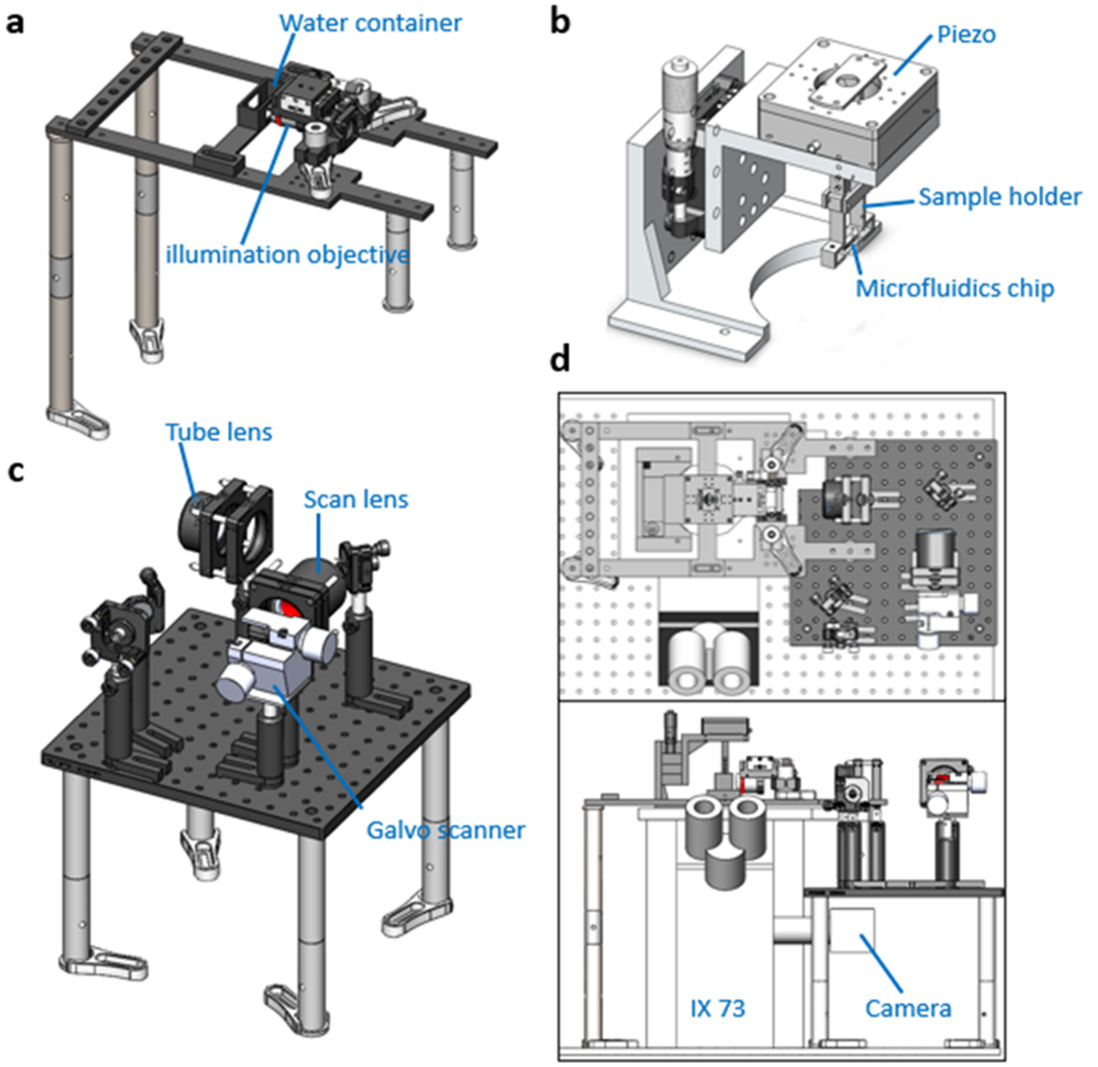


**Figure S2. Design of hardware modules of the DO-DLSM.** (a) The supporting bracket above the microscope. A water container and an illumination objective were mounted on the customized bracket to keep them separated from the two-dimensional mechanical stage of the microscope. (b) The piezo-actuated mechanical module. A step-motor was used for roughly adjusting the height of the piezo. A microfluidics chip was connected to the piezo by a customized sample holder, with both of them being scanned across the light sheet back and forward by the piezo actuator. (c) The plane illumination module. The Lenses and mechanical devices were placed on a 30×30 cm solid aluminum optical breadboard, which was lifted by four pedestal pillars. (d) The top view and front view of the hardware system. The supporting bracket mentioned in (a) and the piezo-actuated mechanical module mentioned in (b) were fixed above the microscope. The plane illumination module was placed on the right of the microscope.

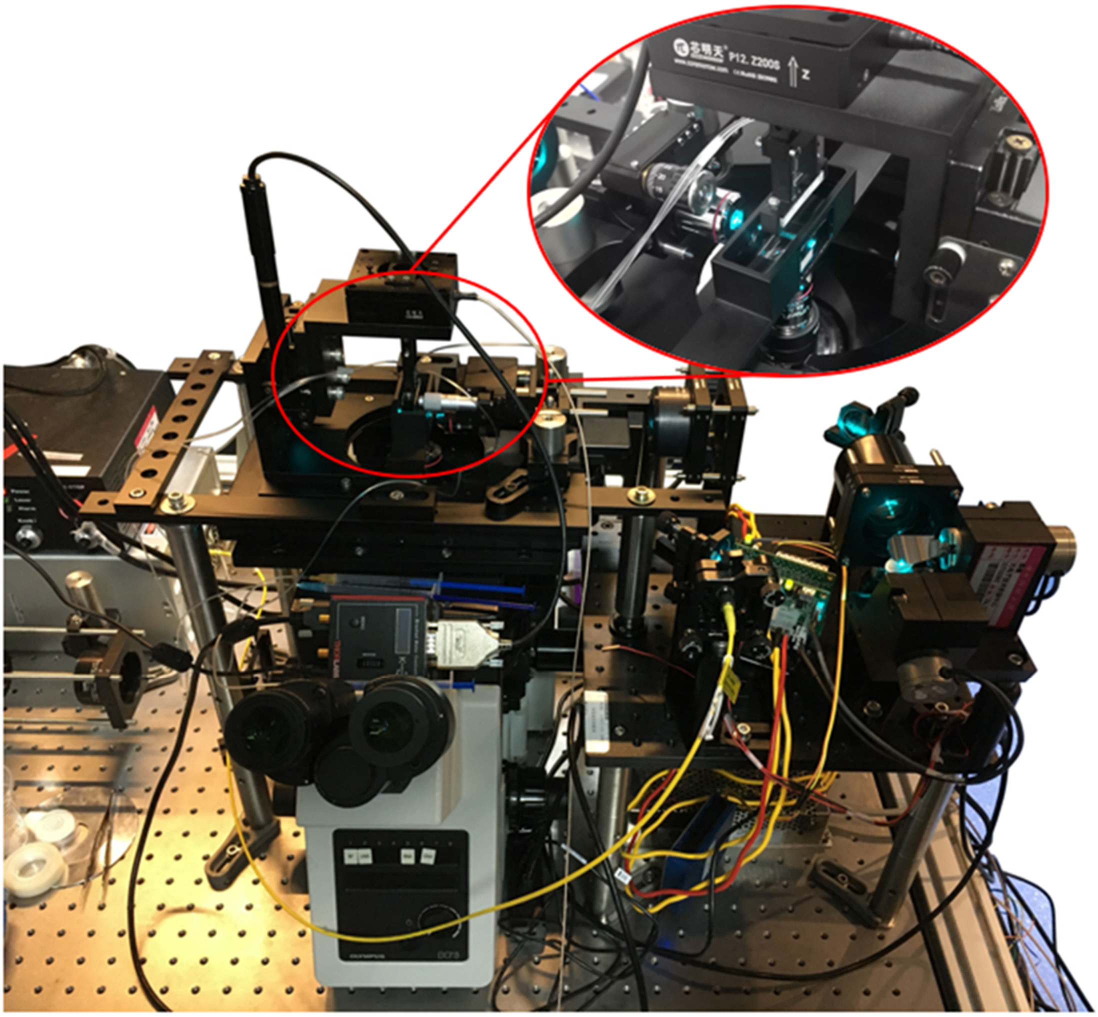

**Figure S3. Photograph of the DO-DSLM system.** The Inset photo shows the microfluidic chip being illuminated by the horizontal light sheet (wavelength, 488 nm).

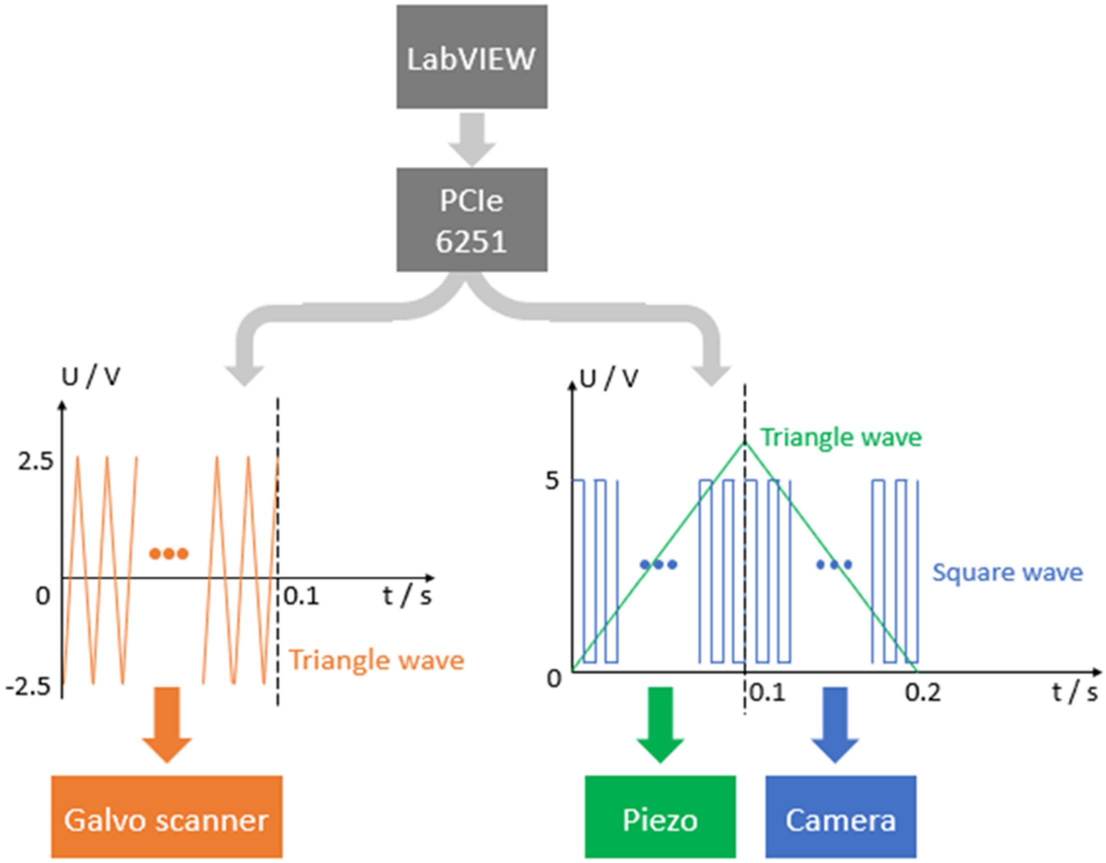


**Figure S4. Signal control in DO-DSLM.** The PCIe 6251 device (NI Instruments) communicated with the computer using customized LabVIEW program. It sent analog and digital signals to the galvo scanner (orange), piezo stage (green) and camera (blue). The galvo scanner received a high frequency triangle wave, with the piezo receiving a low frequency one in the meantime. Next, the camera was trigged by a high frequency square wave. All the signals were arranged in a tightly-controlled time sequence.

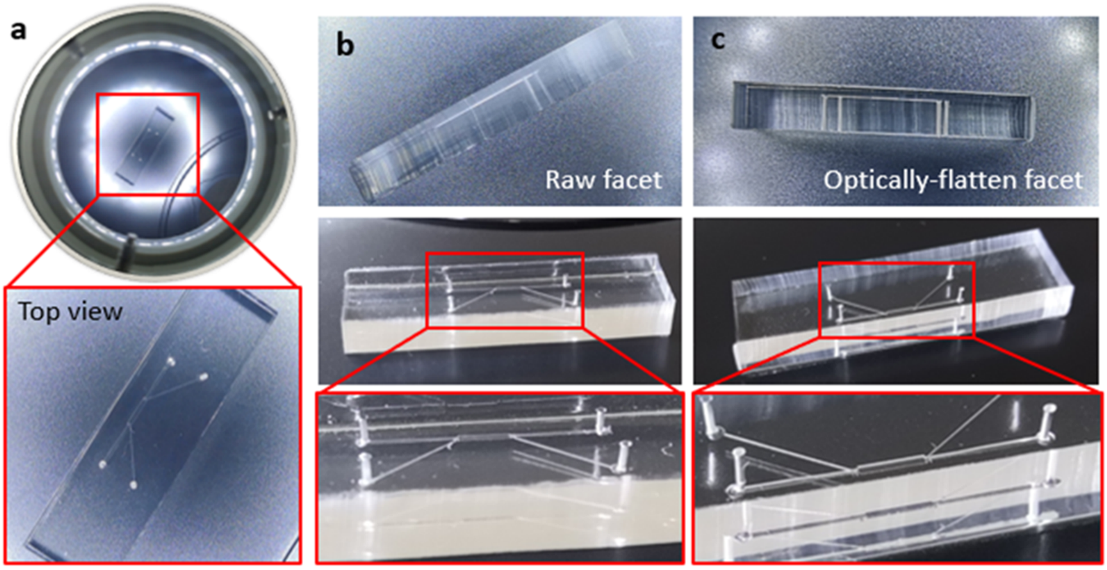


**Figure S5. Side facet flattening of microfluidics chip for horizontal light-sheet illumination.** (a) The photograph of the microfluidics chip from top view. (b) The raw facet of the chip before surface treatment. (c) Optically-flatten facet of the chip after surface treatment.

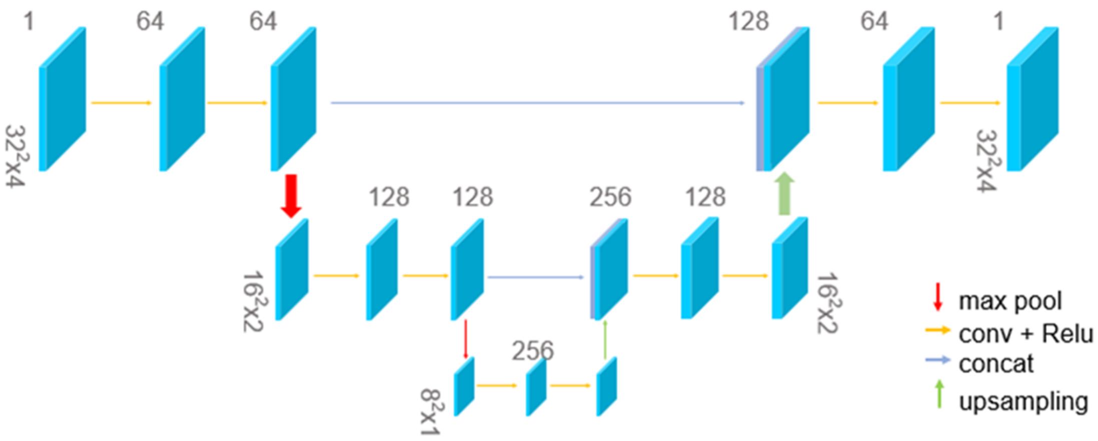


**Figure S6. Architecture of the De-net and Iso-net.** The model contains max-pooling, convolution, concat and upsampling operations, which are represented by red, yellow, blue and green arrows, respectively.

#### Supplementary video

**Video S1.** 3D visualization and quantitative analysis of neural activities in crawling Drosophila larva.

#### Supplementary reference


1 L. Fang, F. Monroe, S. W. Novak, L. Kirk, C. R. Schiavon, S. B. Yu, T. Zhang, M. Wu, K. Kastner, A. A. Latif, Z. Lin, A. Shaw, Y. Kubota, J. Mendenhall, Z. Zhang, G. Pekkurnaz, K. Harris, J. Howard and U. Manor, *Nat. Methods*, 2021, **18**, 406–416.
